# *Apis mellifera* filamentous virus from a honey bee gut microbiome survey in Hungary

**DOI:** 10.1101/2023.07.17.549372

**Authors:** Márton Papp, Adrienn Gréta Tóth, László Békési, Róbert Farkas, László Makrai, Gergely Maróti, Norbert Solymosi

## Abstract

In Hungary, as part of a nationwide, climatically balanced survey for a next-generation sequencing-based study of the honey bee (*Apis mellifera*) gut microbiome, repeated sampling was carried out during the honey production season (March and May 2019). Among other findings, the presence of *Apis mellifera* filamentous virus (AmFV) was detected in all samples, some at very high levels. AmFV-derived reads were more abundant in the March samples than in the May samples. In March, a higher abundance of AmFV-originated reads was identified in samples collected from warmer areas compared to those collected from cooler areas. A lower proportion of AmFV-derived reads were identified in samples collected in March from the wetter areas than those collected from the drier areas. AmFV-read abundance in samples collected in May showed no significant differences between groups based on either environmental temperature or precipitation. The AmFV abundance correlated negatively with *Bartonella apihabitans, Bartonella choladocola*, and positively with *Frischella perrara, Gilliamella apicola, Gilliamella* sp. ESL0443, *Lactobacillus apis, Lactobacillus kullabergensis, Lactobacillus* sp. IBH004. De novo metagenome assembly of four samples resulted in almost the complete AmFV genome. According to phylogenetic analysis based on DNA polymerase, the Hungarian strains are closest to the strain CH-05 isolated in Switzerland.

## Introduction

Honey bees (*Apis mellifera*) are important pollinators with high economic value and ecosystem importance,^1–3^and are exposed to confined environments, and several factors threaten their health, including various pathogens, parasites, and chemicals used as pesticides in agriculture^4–6^. The global decline of this important pollinator poses a threat to food security and biodiversity conservation^7^. The composition of the honey bee’s normal or altered microbiota, for which the available knowledge is limited, may also affect their body function. Although there are studies on honey bee gut microbiota^8–10^, there is little evidence on the environmental factors that influence it. Some results show that seasonal and environmental factors can influence the composition of the gut bacteriome in honeybees.^11–15^. The honey bee gut bacteriome has long been known to be composed of a few core bacterial species^11,16,17^. However, beyond the bacteriome, the viral composition of the microbiome is increasingly gaining more attention in bees^18^, as in humans^19,20^. Most of these studies understandably focus on bacteriophages as they play an essential role in shaping the composition of the bacteriome^18–21^. However, the viruses of the honey bee itself might be just as important. Especially considering those found in the gut and feces, which can contribute to their spread^22–24^. Many of these viruses are important pathogens^22,23^, while others, such as the *Apis mellifera* Filamentous Virus (AmFV), are little to not pathogenic to bees^25,26^. AmFV is the most significant DNA virus of the honey bee^25,26^and for a long time was the only known one^27^. This property of the virus is important because it allows us to investigate the relationship between the bacteriome and AmFV in metagenomic studies targeting bacteria. Despite the fact that the role of AmFV in various diseases is still uncertain, a better understanding of its ecology can bring us closer to understanding its role.

In 2019, a nationwide, climatically balanced survey was conducted in Hungary to investigate changes in the gut microbiome of honey bees based on next-generation sequencing (NGS). The bacteriome results of the study were published by Papp et al. (2022).^15^ In the course of the analyses, we found a large number of short reads originating from AmFV. In this paper, we present the results of a detailed analysis of the AmFV sequences found in the survey.

## Methods

### Sample collection and preparation

Details of the design and conduct of the sampling can be found in the materials and methods section of Papp et al. (2022)^15^, here, we summarise only the methodological details necessary for interpreting the results. In 2019, a country-wide sample collection was performed twice during the honey-producing season, at its onset (March) and the peak (May). A total of 20 sampled apiaries (Fig 1) were selected to obtain a representative sample according to their climatic environment. The climatic environment was characterized by yearly growing degree days (GDD) and the yearly total precipitation. We defined the two categories for our environmental variables as cooler-warmer and less-more for GDD and precipitation, respectively. From individuals collected from three families per apiary, the gastrointestinal tracts of 10 healthy workers per family were removed for sequencing. Paired-end reads were generated from pooled samples per apiary using an Illumina NextSeq sequencer.

**Figure 1.**
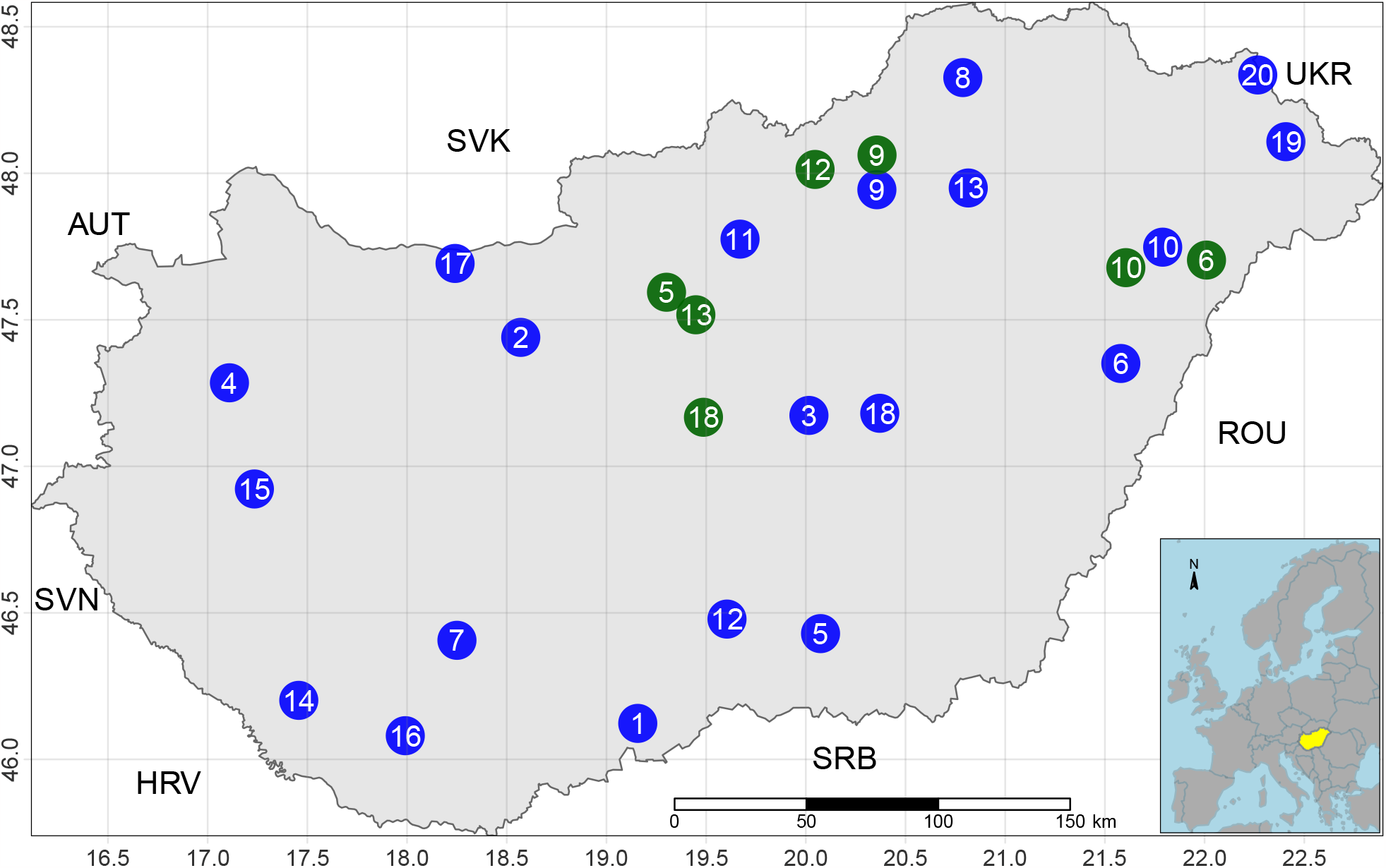
Sampling points in Hungary. The green dots indicate migrating apiaries in May. The blue dots indicate migratory apiaries at the time of sampling in March, as well as non-migratory apiaries. The inset map shows the study region, Hungary in Europe, colored yellow. Neighboring countries are presented by ISO 3 character codes: Austria (AUT), Croatia (HRV), Romania (ROU), Serbia (SRB), Slovakia (SVK), Slovenia (SVN), Ukraine (UKR).

### Bioinformatic and statistical analysis

Quality-based filtering and trimming were performed by Adapterremoval^28^ using 20 as the quality threshold and only retaining reads longer than 50 bp. The remaining reads were taxonomically classified using Kraken2 (k=35)^29^ with the NCBI non-redundant nucleotide database^30^. The taxon classification data was managed using functions of package phyloseq^31^ and microbiome^32^. The abundance differences were analyzed by the DESeq2 package.^33^ Analyzing the seasonal effect, a mixed-effect model was applied to handle the repeated measures by apiary as a random factor. The SparCC correlation coefficient quantified the relationship between the relative abundances of core microbiome species and AmFV.^34,35^The core microbiome was defined with a relative abundance on species-level above 0.5% in at least one of the samples. The statistical tests were two-sided, and p-values less than 0.05 were considered significant. The cleaned reads were aligned to the AmFV genome (KR819915.2) by Bowtie2^36^ with the very-sensitive-local setting. De novo assembly was carried out using MEGAHIT (v1.2.9)^37^, polishing of the contigs was performed with POLCA (v4.1.0)^38^, and scaffolds were created by RagTag (v2.1.0)^39^ using the AmFV genome (KR819915.2).^25^ The average nucleotide identity (ANI) of scaffolds compared to the genome KR819915.2 was estimated by pyani (v0.2.12).^40^ For the genome annotation Prokka (v1.14.6)^41^ was used guided by the genome KR819915.2. Predicted protein homology analysis was performed using the NCBI BLASTP (v2.14.0)^42^ algorithm with a minimum e-value of 1.0e-5 on two reference genomes (KR819915.2, OK392616.1). Phylogenetic analysis was performed based on the amino acid sequences of the DNA polymerase gene. The phylogenetic tree was constructed^43^ based on multiple sequence alignment by MAFFT (v7.490).^44^ The best substitution model was selected by functions of phangorn package^45^ based on the Bayesian information criterion. The generated neighbor-joining tree was optimized by the maximum likelihood method. Bootstrap values were produced by 100 iterations. All data processing and plotting were done in the R-environment.^46^

## Results

The shotgun sequencing generated paired-end read counts of samples are ranging between 311,931 and 546,924, with a mean of 413,629. The OTU table, created by Kraken2 taxonomic classification, contained counts of samples ranging between 175,576 and 314,586 with a median of 262,292. The minimum, maximum, and median read counts of the samples assigned as viral-originated were 443, 72,010, and 1,074, respectively.

The viral hits were dominated by reads matching the genome of AmFV. All of the samples contained reads from this species, and their relative abundance per sample is summarised in Fig 2. AmFV-derived reads were more abundant in the March samples than in the May samples (fold change (FC): 5.53, 95%CI: 2.38-12.84, p<0.001). In March, a higher abundance of AmFV-originated reads was identified in samples collected from warmer areas compared to those collected from cooler areas (FC: 26.05, 95%CI: 7.31-92.81, p<0.001). A lower proportion of AmFV-derived reads were identified in samples collected in March from the wetter areas than in those collected from the drier areas (FC: 0.33, 95%CI: 0.13-0.8, p=0.014). The level of AmFV-reads found in samples collected in May showed no significant differences between groups based on either environmental temperature or precipitation (FC: 1.44, 95%CI: 0.61-3.39, p=0.40; FC: 1.85, 95%CI: 0.78-4.37, p=0.16).

**Figure 2.**
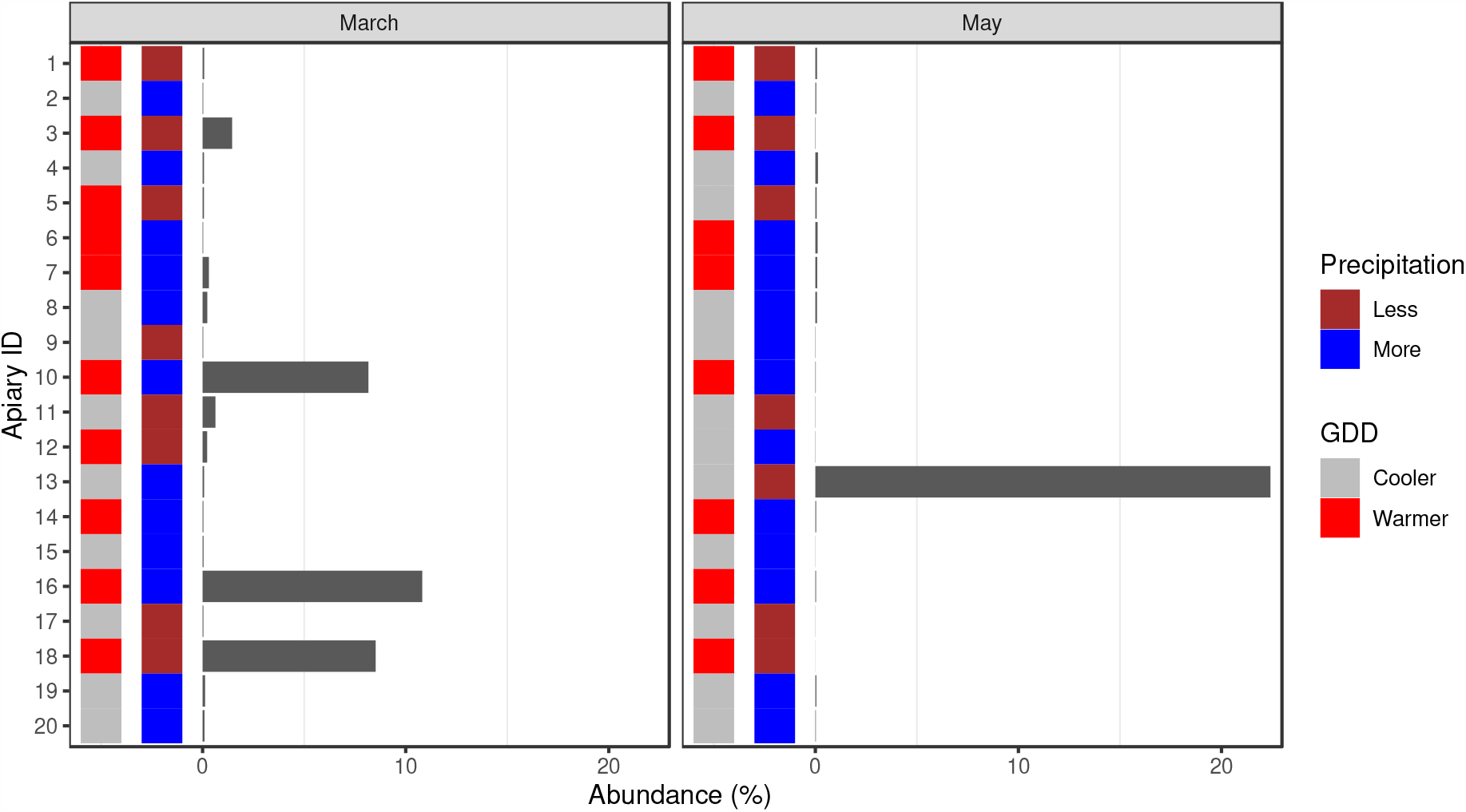
Relative abundance of *Apis mellifera* filamentous virus originated reads for the first (March) and second (May) sampling. The environmental condition, growing degree-day (GDD), and precipitation categories of sampling sites are also marked.

From the core microbiome species *Bartonella apihabitans* (r=-0.384, p<0.001), *Bartonella choladocola* (r=-0.341, p<0.001), *Frischella perrara* (r=0.549, p<0.001), *Gilliamella apicola* (r=0.513, p<0.001), *Gilliamella* sp. ESL0443 (r=0.502, p=0.004), *Lactobacillus apis* (r=0.532, p<0.001), *Lactobacillus kullabergensis* (r=0.515, p=0.002), *Lactobacillus* sp. IBH004 (r=0.399, p<0.001), *Snodgrassella alvi* (r=0.485, p=0.028) showed a significant correlation with AmFV.

De novo assembly of four samples (apiary ID 10, 16, 18 in March and 13 in May) resulted in almost the complete AmFV genome. For these samples, Table 1 shows the coverage and depth of the reads over the reference genome (KR819915.2) and statistics on the scaffolds and the ORFs identified within them.

**Table 1.** Alignment and assembly statistics. Columns 2 and 3 describe the coverage of reads on the reference genome and average depth. The lengths of the scaffolds created by the de novo assembly and the proportion of gaps within them are shown in columns 5 and 6. Column 7 shows the average nucleotide identity (ANI) of scaffolds estimated for the reference genome (KR819915.2). The last two columns show how many ORFs were predicted in the scaffolds generated and how many of these ORFs were predicted to have a protein product that matched a CDS product of the reference genome.

| Apiary ID / month | Reads coverage (%) | depth (x) | MAG GenBank ID | length (bp) | gaps (%) | ANI (%) | ORF predicted | matched |
| --- | --- | --- | --- | --- | --- | --- | --- | --- |
| 10 / March | 98.8 | 8 | OR644611.1 | 504,639 | 2.09 | 97.78 | 153 | 70 |
| 16 / March | 99.1 | 13 | OR644609.1 | 502,090 | 0.84 | 97.78 | 292 | 139 |
| 18 / March | 99.0 | 10 | OR644610.1 | 501,579 | 0.74 | 97.78 | 231 | 119 |
| 13 / May | 99.6 | 31 | OR270109.1 | 499,660 | 0.60 | 97.75 | 286 | 139 |

Table 2 summarises the predicted proteins in the scaffolds generated from our samples that can be linked to products with predicted functions in the KR819915.2 and/or OK392616.1 genomes. Figure 3 shows the phylogenetic tree based on the amino acid sequences of the DNA polymerase gene with the best substitution model JTT+I.

**Table 2.** ORF homologs correspond to genes with predicted functions in the KR819915.2 and/or OK392616.1 genomes. The last four columns show the coverage and sequence identity of the product of the ORFs predicted in our scaffolds that match the product of the reference genome (KR819915.2) with the highest similarity.

| KR819915.2<br>Accession ID | product | OK392616.1<br>Accession ID | product | ORF coverage% / identity% in scaffolds |  |  |  |
| --- | --- | --- | --- | --- | --- | --- | --- |
|  |  |  |  | OR270109.1 | OR644609.1 | OR644610.1 | OR644611.1 |
| AKY03074.1 | zinc-dependent metalloprotease | UQL06506.1 | hypothetical protein | 41.5 / 99.3 | 100.9 / 97.0 | 100.9 / 97.4 | 41.5 / 99.5 |
| AKY03075.1 | hypothetical protein | UQL06507.1 | PK | 100.0 / 99.6 | 100.0 / 98.8 | 100.0 / 99.9 | 100.2 / 98.8 |
| AKY03077.1 | hypothetical protein | UQL06509.1 | BRO-1 | 92.8 / 99.4 | 99.4 / 99.4 | 99.4 / 99.4 | 99.4 / 99.4 |
| AKY03078.1 | hypothetical protein | UQL06510.1 | PARP | 100.2 / 96.8 | 100.2 / 96.6 | 100.0 / 99.8 | 99.3 / 96.2 |
| AKY03080.1 | kinesin motor domain | UQL06512.1 | hypothetical protein | 100.0 / 99.1 | 100.0 / 99.1 | 100.0 / 99.3 | 100.0 / 99.0 |
| AKY03085.1 | BRO domain | UQL06516.1 | BRO-2 | 100.4 / 90.3 | 100.0 / 91.0 | 100.2 / 90.7 | 99.8 / 90.4 |
| AKY03088.1 | hypothetical protein | UQL06519.1 | putative BTB/POZ domain-containing |  |  |  |  |
| AKY03092.1 | AAA+-type ATPase | UQL06523.1 | AAA family ATPase | 100.0 / 99.5 | 100.0 / 99.5 | 100.0 / 99.5 | 100.0 / 99.5 |
| AKY03096.1 | thymidylate synthase | UQL06526.1 | hypothetical protein | 100.0 / 99.3 | 100.7 / 98.4 | 100.7 / 98.5 | 100.0 / 98.9 |
| AKY03097.1 | hypothetical protein | UQL06527.1 | thymidylate synthase |  |  |  |  |
| AKY03103.1 | hypothetical protein | UQL06531.1 | serine protease inhibitor | 95.0 / 94.0 | 96.2 / 95.6 | 38.7 / 99.5 | 95.8 / 95.0 |
| AKY03111.1 | tyrosine recombinase | UQL06534.1 | integrase, partial | 100.0 / 99.5 | 100.0 / 99.6 | 100.0 / 98.5 | 100.0 / 98.9 |
| AKY03112.1 | hypothetical protein | UQL06535.1 | myristoylated membrane | 99.1 / 98.3 | 99.1 / 98.2 | 99.3 / 98.7 | 99.4 / 98.2 |
| AKY03126.1 | hypothetical protein | UQL06546.1 | PIF-5 | 100.6 / 99.1 | 100.6 / 99.1 | 100.6 / 99.1 | 100.6 / 98.8 |
| AKY03129.2 | hypothetical protein | UQL06549.1 | PIF-1 | 105.2 / 99.6 | 100.4 / 99.5 | 100.4 / 99.1 | 30.3 / 99.6 |
| AKY03137.1 | hypothetical protein | UQL06555.1 | putative RING finger protein | 100.0 / 100.0 | 100.0 / 100.0 | 100.0 / 99.7 | 100.0 / 100.0 |
| AKY03138.1 | hypothetical protein | UQL06556.1 | BRO-3 |  | 100.0 / 100.0 | 100.0 / 99.6 | 100.0 / 99.6 |
| AKY03143.1 | DNA polymerase B | UQL06561.1 | DNA polymerase family B | 99.5 / 98.5 | 99.5 / 99.0 | 7.4 / 99.3 | 99.4 / 99.0 |
| AKY03144.1 | hypothetical protein | UQL06562.1 | BRO-4 | 100.0 / 99.8 | 100.0 / 99.8 | 100.0 / 99.8 | 100.0 / 99.8 |
| AKY03146.1 | hypothetical protein | UQL06564.1 | PIF-0 | 99.7 / 99.7 | 99.7 / 99.4 | 99.7 / 99.6 | 99.7 / 99.7 |
| AKY03149.1 | hypothetical protein | UQL06567.1 | RING finger | 98.6 / 97.7 | 100.0 / 100.0 | 99.3 / 98.9 | 100.0 / 99.3 |
| AKY03151.1 | hypothetical protein | UQL06569.1 | HZV 115-like protein | 100.0 / 99.8 | 100.0 / 99.8 | 100.0 / 99.8 | 100.0 / 99.8 |
| AKY03157.1 | pif domain | UQL06575.1 | PIF-3 | 100.0 / 100.0 | 100.0 / 100.0 | 100.0 / 100.0 | 100.0 / 100.0 |
| AKY03164.1 | similar to DNA ligase I | UQL06580.1 | DNA ligase 3 | 99.2 / 98.8 | 99.6 / 99.1 | 99.6 / 99.2 | 99.2 / 98.6 |
| AKY03169.1 | hypothetical protein | UQL06585.1 | PIF-2 | 100.0 / 99.8 | 100.0 / 100.0 | 100.0 / 100.0 | 100.0 / 100.0 |
| AKY03170.1 | similar to bacterial gamma-glutamyltranspeptidase | UQL06586.1 | SEA domain-containing protein Gamma-glutamyltranspeptidase | 99.1 / 97.8 | 99.1 / 98.5 | 99.1 / 98.6 | 98.2 / 97.6 |
| AKY03175.1 | BRO domain | UQL06591.1 | BRO-5 | 96.5 / 93.7 | 96.5 / 94.1 | 53.5 / 52.3 |  |
| AKY03177.1 | hypothetical protein | UQL06593.1 | BRO-6 | 102.1 / 61.1 | 102.1 / 60.3 | 56.5 / 66.3 |  |
| AKY03179.2 | hypothetical protein | UQL06595.1 | BRO-7 | 185.1 / 98.1 | 184.0 / 96.7 | 138.7 / 96.7 |  |
| AKY03180.1 | hypothetical protein | UQL06596.1 | BRO-8 | 100.0 / 98.9 | 100.0 / 97.0 | 100.0 / 98.6 |  |
| AKY03183.1 | ribonucleotide reductase, large subunit | UQL06599.1 | ribonucleotide reductase 1 | 100.0 / 99.9 | 56.6 / 100.0 | 100.0 / 99.9 |  |
| AKY03191.1 | similar to RecB exonuclease | UQL06606.1 | hypothetical protein | 100.0 / 99.8 | 100.0 / 99.4 | 100.0 / 99.8 |  |
| AKY03202.1 | BRO domain | UQL06616.1 | BRO-9 | 100.0 / 100.0 | 100.0 / 99.4 | 100.0 / 100.0 |  |
| AKY03226.1 | hypothetical protein | UQL06638.1 | PIF-4 | 40.9 / 100.0 | 100.0 / 100.0 | 100.0 / 100.0 |  |
| AKY03237.1 | hypothetical protein | UQL06646.1 | MdSGHV070 | 99.7 / 95.3 | 99.8 / 94.8 | 98.7 / 94.6 |  |
| AKY03262.2 | hypothetical protein | UQL06666.1 | chitin-binding type-4 domain-containing protein | 109.5 / 97.5 | 110.0 / 97.6 |  |  |
| AKY03304.1 | similar to bacterial phosphatidylinositol-specific phospholipase C, catalytic domain | UQL06696.1 | phospholipase C | 100.0 / 100.0 | 100.0 / 100.0 |  |  |
| WKY35419.1 | ribonucleotide reductase, small subunit | UQL06680.1 | ribonucleotide reductase 2 | 99.7 / 98.7 | 89.6 / 97.7 |  |  |

**Figure 3.**
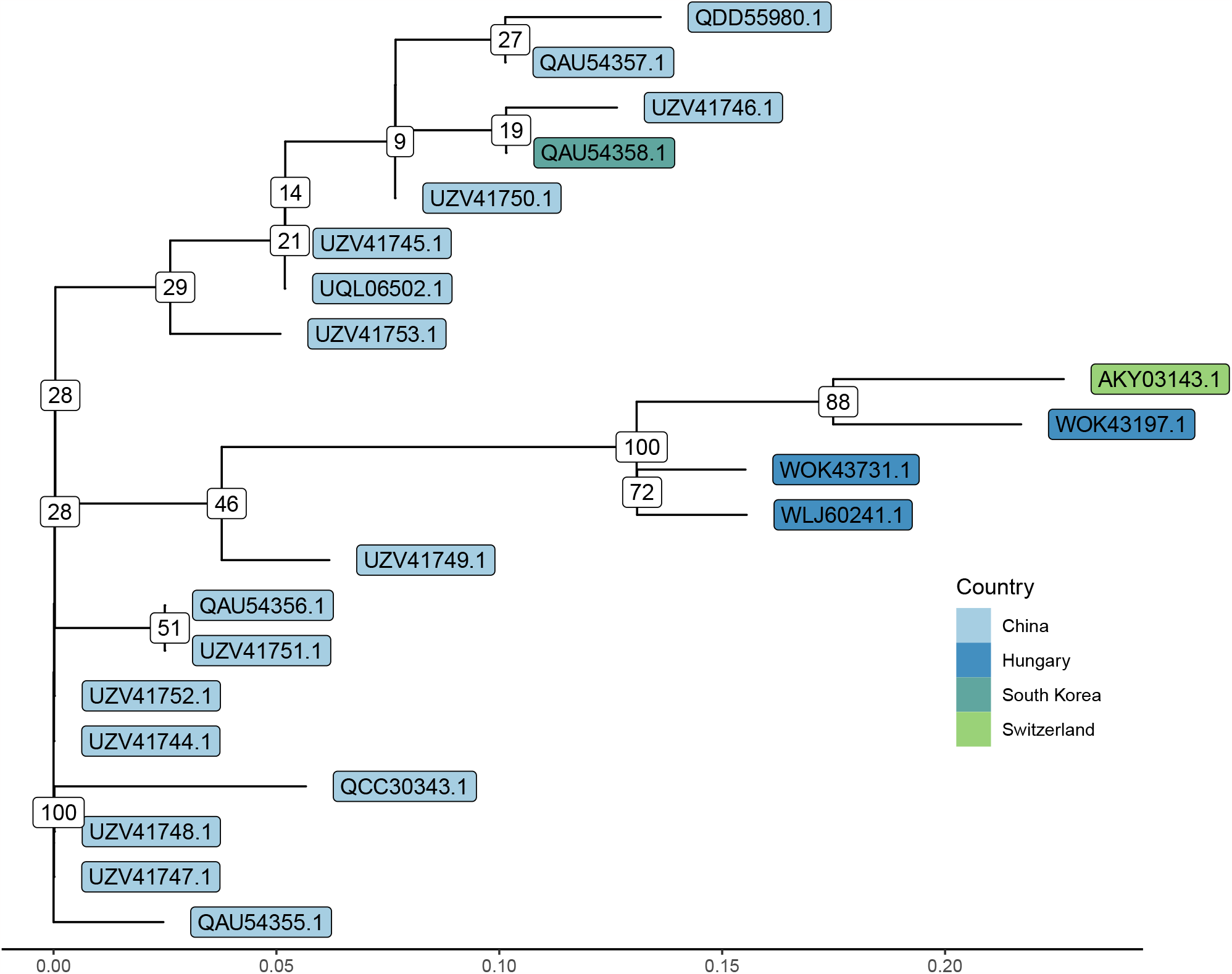
Phylogenetic tree based on DNA-polymerase amino acid sequences. The genes WOK43731.1, WOK43197.1 and WLJ60241.1 were assembled from the samples of apiary 10 and 16 in March and apiary 13 in May, respectively. Most of the sequences derived from *Apis mellifera*, with some from other hosts: *Apis andreniformis* (QCC30343.1), *Apis cerana* (UZV41744.1), *Apis cerana cerana* (QAU54355.1), bumblebee (QAU54357.1), *Galleria mellonella* (QDD55980.1).

## Discussion

Even though samples presented in this study were taken from healthy bee specimens, AmFV was detected in all samples. This is in line with our current knowledge of the virus, according to which it is only pathogenic in acute cases and/or if the bee colony is under stress.^26,47,48^The virus is otherwise commonly prevalent, or even endemic in bee colonies, and besides its very likely oral-fecal transmission route, can possibly be spread transovarially from the queen to the workers.^25^

In contrast to other studies, we have observed a decline in AmFV abundances as the honey-producing season advanced. Bailey et al. observed an increasing prevalence of infected colonies from the beginning of autumn until the end of spring, which was followed by a steep decline^49^. Similarly, Hartmann and colleagues observed a significant difference in AmFV loads between autumn and springtime^26^. It is possible that regional climatic differences might influence the dynamics of viral loads. Further, large-scale surveys are necessary, however, to confirm such patterns.

Furthermore, the relationship of AmFV with the other components of the microbiome can provide us useful information. We found a significant positive correlation with AmFV for several bacteria. In some of these, the abundance of the bacterial species also decreased as the season progressed, as we have observed previously for *L. apis* and *L. kullabergensis*^15^. For other species, such as *F. perrara, G. apicola* and *S. alvi*, we have not observed a seasonal pattern in our previous analysis^15^. Among these, *F. perrara* is especially of interest, which is known to stimulate the immune system of the honey bees strongly^50^. This species, as a potential opportunistic pathogen^51^, may act as a stressor on the bees, thus explaining the observed positive correlation with AmFV. Significant negative correlation was only found for two newly described *Bartonella* species, *B. apihabitans* and *B. choladocola*^52^. Regarding the only previously known *Bartonella* species in bees, *B. apis*, our previous analysis of the same samples found an increase in abundance of this bacterial species with the progression of the season^15^. This might suggest that even these two newly described species may have shown similar changes, which may explain the negative correlation.

The reason for the seasonal peaks of AmFV or the association between the virus and other members of the honey bee microbiome is yet unknown. Furthermore, the effect of annually higher AmFV abundances on the immune system of bees needs to be determined. Moreover, other studies found correlations in the number of AmFv and other RNA viruses, such as Deformed Wing Virus (DWV) and Black Queen Cell Virus (BQCV)^26,53^, indicating the benefit that could be achieved if the microbiome could be analyzed as a whole.

Our results confirm the high prevalence of the virus, as previous studies suggested^26,49^. There were, however, 3 March samples (sample numbers 10, 16, and 18) in which we found an exceptionally high abundance of AmFV. This may be somewhat related to our observations on seasonality, as the virus was not present in such high numbers in the corresponding May samples. Accordingly, the abundance of the virus decreased as the season progressed. In contrast, however, sample 13 was dominated by AmFV in May and, therefore, may be of further interest. The spike in virus abundance may indicate the presence of some stressor on the colony, which has not yet caused observable symptoms in the animals. Bee viruses have been described, for example, to increase in abundance in response to certain pesticides in a concentration-dependent manner^54,55^, which is thought to be due to the immunomodulatory effect of pesticides^54^ If indeed the emergence of stressors could be associated with such a fluctuation in viral abundance, it could also suggest the use of the virus as a health indicator and contribute to the diagnosis of various complex disease processes.

Based on the phylogenetic analysis, the sample from apiary 16, collected in March from the southwest of Hungary, had the highest similarity to the Swiss reference genome. Both other samples (apiary ID 10 March and 13 May) that contained the DNA-polymerase gene in full length were derived from Eastern Hungary. Even though sample 13 from May is relatively closer to the west of the country due to colony migration, it is originally from Eastern Hungary, and the permanent beekeeping premises are 76 air kilometers away from March sampling point 10. Accordingly, it can be supposed that the viral strain from sample 13 collected in May has already been present at the overwintering location and migrated to the May sampling point. It can be assumed that the mediating effect of the imports of honey and propolis from Hungary to Switzerland is responsible for the close genetic relationship of the Hungarian strains of AmFV.

More than one hundred ORFs were identified in each of four of our scaffolds that did not show any similarity to the reference genome with the homology assumptions. In a previous study, Yang and colleagues (2022)^56^ predicted the functionality of newly detected ORFs from AmFV isolates. As our sequences were of metagenomic origins, the predicted products may derive from organisms other than AmFV despite our sequences’ high similarity to the reference sequence. Accordingly, we have presented only those products that appeared in any of the two whole genomes.

In the present study, we report 4 new assemblies of the *Apis mellifera* filamentous virus from Hungary. Our results may provide deeper insights into the genome organization of this virus. Furthermore, our results suggest a seasonal trend of the virus abundance, i.e., there is a decrease in it in the gastrointestinal tract of bees as the production season progresses. The high abundance of the virus in some bee colonies and its association with certain members of the microbiome, in particular *Frischella perrara*, may suggest a link between the virus and various stressors. It may, therefore, be important either as an agent of complex disease processes or as an indicator of the health status of the hive.

## Acknowledgments

In memory of Rajnald András Köveshegyi OCist. We would like to say thanks to the beekeepers for giving us their indispensable help. It has also received funding from the European Union’s Horizon 2020 research and innovation program under Grant Agreement No. 874735 (VEO). GM received support from the Hungarian Academy of Sciences through the Lendület-Programme (LP2020-5/2020). Supported by the ÚNKP-22-3-II. New National Excellence program of the Ministry for Culture and Innovation from the source of the National Research, Development and Innovation Fund.

## Author contributions statement

NS takes responsibility for the integrity of the data and the accuracy of the data analysis. LB, LM, NS, and RF conceived the concept of the study. GM, LB, LM, NS, and RF performed sample collection and procedures. AGT, MP, and NS participated in the bioinformatic and statistical analysis. AGT, MP, and NS participated in the drafting of the manuscript. AGT, GM, LB, LM, MP, NS, and RF carried out the critical revision of the manuscript for important intellectual content. All authors read and approved the final manuscript.

## Additional information

### Availability of data and material

The short read data of sample data are publicly available and can be accessed through the PRJNA685398 from the NCBI Sequence Read Archive (SRA).

### Ethics approval and consent to participate

Not applicable.

### Consent for publication

Not applicable.

## References

1. Ványi, G. Á., Csapó, Z. & Kárpáti, L. Externality effects of honey production. Appl. Stud. Agribusiness Commer. 6, 63–67, DOI: 10.19041/APSTRACT/2012/1-2/8 (2012).

2. Hristov, P., Neov, B., Shumkova, R. & Palova, N. Significance of Apoidea as main pollinators. Ecological and economic impact and implications for human nutrition. Diversity 12, 280, DOI: 10.3390/d12070280 (2020).

3. Patel, V., Pauli, N., Biggs, E., Barbour, L. & Boruff, B. Why bees are critical for achieving sustainable development. Ambio 50, 49–59, DOI: 10.1007/s13280-020-01333-9 (2021).

4. Oldroyd, B. P. What’s killing American honey bees? PLoS Biol. 5, e168, DOI: 10.1371/journal.pbio.0050168 (2007).

5. Barbosa, W. F., Smagghe, G. & Guedes, R. N. C. Pesticides and reduced-risk insecticides, native bees and pantropical stingless bees: pitfalls and perspectives. Pest Manag. Sci. 71, 1049–1053, DOI: 10.1002/ps.4025 (2015).

6. Morawetz, L. et al. Health status of honey bee colonies (Apis mellifera) and disease-related risk factors for colony losses in Austria. PloS ONE 14, e0219293, DOI: 10.1371/journal.pone.0219293 (2019).

7. Potts, S. G. et al. Global pollinator declines: trends, impacts and drivers. Trends Ecol. & Evol. 25, 345–353, DOI: 10.1016/j.tree.2010.01.007 (2010).

8. Moran, N. A., Hansen, A. K., Powell, J. E. & Sabree, Z. L. Distinctive gut microbiota of honey bees assessed using deep sampling from individual worker bees. PLoS ONE 7, DOI: 10.1371/journal.pone.0036393 (2012).

9. Anderson, K. E. et al. The queen’s gut refines with age: longevity phenotypes in a social insect model. Microbiome 6, 1–16, DOI: 10.1186/s40168-018-0489-1 (2018).

10. Regan, T. et al. Characterisation of the British honey bee metagenome. Nat. Commun. 9, 1–13, DOI: 10.1038/s41467-018-07426-0 (2018).

11. Subotic, S. et al. Honey bee microbiome associated with different hive and sample types over a honey production season. PloS ONE 14, e0223834, DOI: 10.1371/journal.pone.0223834 (2019).

12. Ludvigsen, J. et al. Shifts in the midgut/pyloric microbiota composition within a honey bee apiary throughout a season. Microbes Environ. 30, 235–244, DOI: 10.1264/jsme2.ME15019 (2015).

13. Corby-Harris, V., Maes, P. & Anderson, K. E. The bacterial communities associated with honey bee (Apis mellifera) foragers. PloS ONE 9, e95056, DOI: 10.1371/journal.pone.0095056 (2014).

14. Kešnerová, L. et al. Gut microbiota structure differs between honeybees in winter and summer. The ISME J. 14, 801–814, DOI: 10.1038/s41396-019-0568-8 (2020).

15. Papp, M. et al. Natural diversity of the honey bee (Apis mellifera) gut bacteriome in various climatic and seasonal states. PloS ONE 17, e0273844, DOI: 10.1371/journal.pone.0273844 (2022).

16. Kwong, W. K. & Moran, N. A. Gut microbial communities of social bees. Nat. Rev. Microbiol. 14, 374–384, DOI: 10.1038/nrmicro.2016.43 (2016).

17. Jones, J. C. et al. Gut microbiota composition is associated with environmental landscape in honey bees. Ecol. Evol. 8, 441–451, DOI: 10.1002/ece3.3597 (2018).

18. Bonilla-Rosso, G., Steiner, T., Wichmann, F., Bexkens, E. & Engel, P. Honey bees harbor a diverse gut virome engaging in nested strain-level interactions with the microbiota. PNAS 117, 7355–7362, 10.1073/pnas.2000228117 (2020).

19. Columpsi, P. et al. Beyond the gut bacterial microbiota: The gut virome. J. Med. Virol. 88, 1467–1472, DOI: 10.1002/jmv.24508 (2016).

20. Garmaeva, S. et al. Studying the gut virome in the metagenomic era: challenges and perspectives. BMC Biol. 17, 1–14, DOI: 10.1186/s12915-019-0704-y (2019).

21. Bueren, E. K. et al. Characterization of prophages in bacterial genomes from the honey bee (Apis mellifera) gut microbiome. PeerJ 11, DOI: 10.7717/peerj.15383 (2023).

22. Chen, Y., Evans, J. & Feldlaufer, M. Horizontal and vertical transmission of viruses in the honey bee, Apis mellifera. J. Invertebr. Pathol. 92, 152–159, DOI: 10.1016/j.jip.2006.03.010 (2006).

23. Chen, Y. P. & Siede, R. Honey bee viruses. Adv. Virus Res. 70, 33–80, DOI: 10.1016/S0065-3527(07)70002-7 (2007).

24. Daughenbaugh, K. F. et al. Honey bee infecting Lake Sinai viruses. Viruses 7, 3285–3309, DOI: 10.3390/v7062772 (2015).

25. Gauthier, L. et al. The Apis mellifera filamentous virus genome. Viruses 7, 3798–3815, DOI: 10.3390/v7072798 (2015).

26. Hartmann, U., Forsgren, E., Charrière, J.-D., Neumann, P. & Gauthier, L. Dynamics of Apis mellifera filamentous virus (AmFV) infections in honey bees and relationships with other parasites. Viruses 7, 2654–2667, DOI: 10.3390/v7052654 (2015).

27. Kraberger, S. et al. Diverse single-stranded DNA viruses associated with honey bees (Apis mellifera). Infect. Genet. Evol. 71, 179–188, DOI: 10.1016/j.meegid.2019.03.024 (2019).

28. Schubert, M., Lindgreen, S. & Orlando, L. AdapterRemoval v2: rapid adapter trimming, identification, and read merging. BMC Res. Notes 9, 88, DOI: 10.1186/s13104-016-1900-2 (2016).

29. Wood, D. E., Lu, J. & Langmead, B. Improved metagenomic analysis with Kraken 2. Genome Biol. 20, 1–13, DOI: 10.1186/s13059-019-1891-0 (2019).

30. Pruitt, K. D., Tatusova, T. & Maglott, D. R. NCBI reference sequence (RefSeq): a curated non-redundant sequence database of genomes, transcripts and proteins. Nucleic Acids Res. 33, D61–D65, DOI: 10.1093/nar/gkl842 (2007).

31. McMurdie, P. J. & Holmes, S. phyloseq: An R package for reproducible interactive analysis and graphics of microbiome census data. PLoS ONE 8, 1–11, DOI: 10.1371/journal.pone.0061217 (2013).

32. Lahti, L. & Shetty, S. microbiome R package (2012-2019). http://microbiome.github.io.

33. Love, M. I., Huber, W. & Anders, S. Moderated estimation of fold change and dispersion for RNA-seq data with DESeq2. Genome Biol. 15, 1–21, DOI: 10.1186/s13059-014-0550-8 (2014).

34. Friedman, J. & Alm, E. J. Inferring correlation networks from genomic survey data. PLoS Comput. Biol. 8, 1–11, DOI: 10.1371/journal.pcbi.1002687 (2012).

35. Kurtz, Z. D. et al. Sparse and compositionally robust inference of microbial ecological networks. PLoS Comput. Biol. 11, 1–25, DOI: 10.1371/journal.pcbi.1004226 (2015).

36. Langmead, B. & Salzberg, S. L. Fast gapped-read alignment with Bowtie 2. Nat. Methods 9, 357–359, DOI: 10.1038/nmeth.1923 (2012).

37. Li, D., Liu, C.-M., Luo, R., Sadakane, K. & Lam, T.-W. MEGAHIT: an ultra-fast single-node solution for large and complex metagenomics assembly via succinct de Bruijn graph. Bioinformatics 31, 1674–1676, DOI: 10.1093/bioinformatics/btv033 (2015).

38. Zimin, A. V. & Salzberg, S. L. The genome polishing tool POLCA makes fast and accurate corrections in genome assemblies. PLoS Comput. Biol. 16, e1007981, DOI: 10.1371/journal.pcbi.1007981 (2020).

39. Alonge, M. et al. Automated assembly scaffolding using RagTag elevates a new tomato system for high-throughput genome editing. Genome Biol. 23, 258, DOI: 10.1186/s13059-022-02823-7 (2022).

40. Pritchard, L., Glover, R. H., Humphris, S., Elphinstone, J. G. & Toth, I. K. Genomics and taxonomy in diagnostics for food security: soft-rotting enterobacterial plant pathogens. Anal. Methods 8, 12–24, DOI: 10.1039/C5AY02550H (2016).

41. Seemann, T. Prokka: rapid prokaryotic genome annotation. Bioinformatics 30, 2068–2069, DOI: 10.1093/bioinformatics/btu153 (2014).

42. Camacho, C. et al. Blast+: architecture and applications. BMC Bioinforma. 10, 1–9, DOI: 10.1186/1471-2105-10-421 (2009).

43. Yu, G., Smith, D., Zhu, H., Guan, Y. & Lam, T. T.-Y. ggtree: an R package for visualization and annotation of phylogenetic trees with their covariates and other associated data. Methods Ecol. Evol. 8, 28–36, DOI: 10.1111/2041-210X.12628 (2017).

44. Katoh, K. & Standley, D. M. MAFFT multiple sequence alignment software version 7: improvements in performance and usability. Mol. Biol. Evol. 30, 772–780, DOI: 10.1093/molbev/mst010 (2013).

45. Schliep, K., Potts, A. J., Morrison, D. A. & Grimm, G. W. Intertwining phylogenetic trees and networks. Methods Ecol. Evol. 8, 1212–1220, DOI: 10.1111/2041-210X.12760 (2017).

46. R Core Team. R: A Language and environment for statistical computing. R Foundation for Statistical Computing, Vienna, Austria (2023). https://www.R-project.org/.

47. Clark, T. B. A filamentous virus of the honey bee. J. Invertebr. Pathol. 32, 332–340, DOI: 10.1016/0022-2011(78)90197-0 (1978).

48. Federici, B. A., Bideshi, D. K., Tan, Y., Spears, T. & Bigot, Y. Ascoviruses: Superb Manipulators of Apoptosis for Viral Replication and Transmission, 171–196 (Springer Berlin Heidelberg, Berlin, Heidelberg, 2009).

49. Bailey, L., Ball, B. V. & Perry, J. Association of viruses with two protozoal pathogens of the honey bee. Annals Appl. Biol. 103, 13–20, DOI: 10.1111/j.1744-7348.1983.tb02735.x (1983).

50. Emery, O., Schmidt, K. & Engel, P. Immune system stimulation by the gut symbiont Frischella perrara in the honey bee (Apis mellifera). Mol. Ecol. 26, 2576–2590, DOI: 10.1111/mec.14058 (2017).

51. Maes, P. W., Rodrigues, P. A., Oliver, R., Mott, B. M. & Anderson, K. E. Diet-related gut bacterial dysbiosis correlates with impaired development, increased mortality and Nosema disease in the honeybee (Apis mellifera). Mol. Ecol. 25, 5439–5450, DOI: 10.1111/mec.13862 (2016).

52. Liu, Y., Chen, J., Lang, H. & Zheng, H. Bartonella choladocola sp. nov. and Bartonella apihabitans sp. nov., two novel species isolated from honey bee gut. Syst. Appl. Microbiol. 45, 126372, DOI: 10.1016/j.syapm.2022.126372 (2022).

53. Abou Kubaa, R. et al. First detection of black queen cell virus, Varroa destructor macula-like virus, Apis mellifera filamentous virus and Nosema ceranae in Syrian honey bees Apis mellifera syriaca. Bull. Insectology 71, 217–224 (2018).

54. Di Prisco, G. et al. Neonicotinoid clothianidin adversely affects insect immunity and promotes replication of a viral pathogen in honey bees. PNAS 110, 18466–18471, DOI: 10.1073/pnas.1314923110 (2013).

55. Coulon, M. et al. Metabolisation of thiamethoxam (a neonicotinoid pesticide) and interaction with the chronic bee paralysis virus in honeybees. Pesticide Biochem. Physiol. 144, 10–18, DOI: 10.1016/j.pestbp.2017.10.009 (2018).

56. Yang, D. et al. Genomics and proteomics of Apis mellifera filamentous virus isolated from honeybees in China. Virol. Sinica 37, 483–490, DOI: 10.1016/j.virs.2022.02.007 (2022).

